# Cytokinesis and post-abscission midbody remnants are regulated during mammalian brain development

**DOI:** 10.1101/529164

**Authors:** Katrina C. McNeely, Noelle D. Dwyer

## Abstract

Building a brain of the proper size and structure requires neural stem cells (NSCs) to divide with tight temporal and spatial control to produce different daughter cell types in proper numbers and sequence. Mammalian NSCs in the embryonic cortex must maintain their polarized epithelial structure as they undergo both early proliferative divisions and later neurogenic divisions. To do this they undergo a polarized form of cytokinesis at the apical membrane that is not well understood. Here we investigate whether polarized furrowing and abscission in mouse NSCs are regulated differently at earlier and later stages, and in a cytokinesis mutant, *Kif20b*. We developed methods to live image furrow ingression and midbody microtubule abscission in NSCs within cortical explants. We find that polarized furrow ingression occurs at a steady rate and completes in 15 minutes, regardless of developmental age. However, ingression is slowed in a subset of *Kif20b* mutant NSCs. Abscission is usually observed on both midbody flanks, and takes 65-75 minutes to complete. Surprisingly, the process is accelerated in the microcephalic *Kif20b* mutant NSCs. Midbody remnants litter the apical membrane surface, and are more prevalent with early proliferative divisions. These results suggest that abscission is developmentally regulated in NSCs and may influence proper brain growth and structure.

## Introduction

To build a brain of the proper size and structure, neural stem cells (NSCs) must proliferate rapidly to produce billions of daughter cells in a short developmental time window, and generate different daughter cell types at specific times. NSCs are tall, thin cells that are highly polarized, extending radially to contact the pia on the basal side. Their apical membranes (“apical endfeet”) are joined by junctions and form the walls of the lateral ventricles. Their nuclei move within them during the cell cycle, in a process called interkinetic nuclear migration. Nuclei move to the basal side for S-phase and to the apical membrane for M-phase and must carefully regulate the positioning of mitosis and cytokinesis (see Figure 1B). This nuclear movement creates a pseudostratified epithelium as NSCs proliferate. During early development, NSCs perform symmetric proliferative divisions to produce two NSC daughters and expand the stem cell pool. Later, NSCs increase neurogenic divisions, producing neuron daughters that differentiate, migrate basally, and never divide again (1, 2). Errors in these divisions can result in brains that are too small or have abnormal structure (2, 3).How the NSCs accomplish these divisions and control the balance of proliferative and neurogenic daughter fates is a subject of intense study.

**Figure 1:**
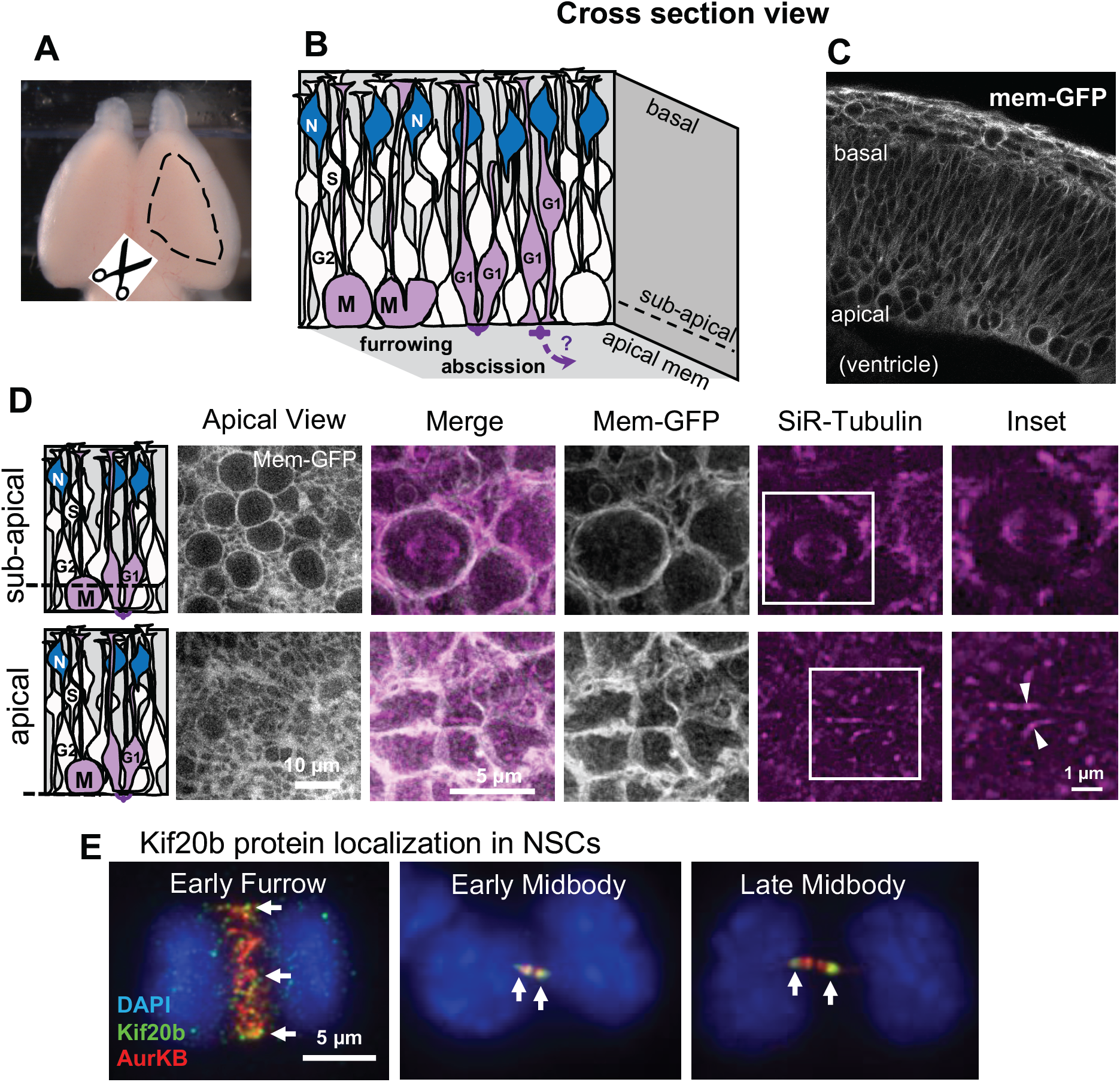
Imaging cytokinesis in neural stem cells of developing cortex. (A) Embryonic mouse brain with dashed outline showing area of cortex dissected for cortical slab preparation. (B) Schematic of the pseudostratified epithelium of the developing cerebral cortex. NSCs undergo mitosis (M), furrowing, and abscission at the apical membrane. NSCs undergo interkinetic nuclear migration. Nuclei move basally for S-phase and apically for mitosis (M). Abscission occurs during G1 phase. N, neurons. (C) Cross section image of E12.5 mouse cortex expressing membrane-GFP shows the dense packing of pseudostratified epithelium. Round cells in mitosis can be seen close to the apical surface. (D) Schematics and images of cortical slabs labeled with membrane-GFP and SiR-tubulin, viewing the apical membrane *en face*, at two different imaging planes. Top row shows the sub-apical plane where larger cell diameters of the mitotic cells with rounded membranes and mitotic spindles are located. Bottom row shows the apical plane where apical endfeet are located, and where the midbody forms and abscission occurs. Arrowheads point to the bulges of two different midbodies. (E) Endogenous Kif20b immunostaining in dissociated embryonic cortical mouse NSCs. Kif20b is enriched on the central spindle of anaphase cells, the flanks of early midbodies, and in the constriction sites of later midbodies (arrows). It is undetectable in *Kif20b* mutant NSCs (26).

As they divide, the NSCs must faithfully segregate genomes and organelles to their daughters, and confer proper daughter fates, while maintaining their polarity and the integrity of the epithelium. To do this, NSCs undergo a polarized form of cytokinesis that is poorly understood: first, the furrow ingresses from basal to apical; then, abscission occurs at the apical membrane. Cleavage is near-perpendicular to the apical membrane (4). While many studies have shown that disruption in cleavage plane can deplete the stem cell pool or disrupt cortical structure (5, 6), the regulation of furrow ingression itself has not been thoroughly addressed.

The basic mechanisms of cytokinetic abscission have been established primarily from studies in single cell models (7-9). After chromosome segregation, the central spindle promotes cleavage furrow ingression and compacts its microtubules into a structure called the midbody, within the intercellular bridge. The midbody contains over 150 proteins that assemble into a central bulge and lateral flanks (10, 11). This organelle mediates abscission, the process of severing the intercellular bridge. Abscission involves both microtubule disassembly and plasma membrane scission at constriction sites on midbody flanks (12, 13). After abscission, the central bulge remains intact and is called the midbody remnant (MBR). Evidence from developing worms and flies, as well as mammalian stem cell lines, suggests that temporal and spatial regulation of abscission can influence daughter cell polarity and fate (14-20). Potentially, MBRs could transmit signals to neighboring cells by surface binding or uptake (21-24). It is unclear whether these simpler systems accurately model abscission dynamics in the developing brain, where polarized stem cells must alter the balance of proliferation and differentiation during development.

Previously, we showed that a loss-of-function mutation of the Kinesin-6 *Kif20b* in mice causes microcephaly (25, 26). Kif20b protein localizes to the central spindle during furrowing, then to midbody flanks and finally to the constriction sites (Figure 1G and (27, 28). This dynamic localization during cytokinesis, suggests that Kif20b could play important roles in furrowing or abscission. We showed that in HeLa cells, Kif20b depletion caused subtle disruptions of furrowing speed, midbody maturation and abscission timing, but does not prevent abscission completion (27). This raised the question of whether Kif20b loss in NSCs would cause more severe defects in cytokinesis. In fixed brains of *Kif20b* mutants, we found no changes in the proportions or positions of mitotic or S-phase NSCs (26). Cleavage furrow angles were not different than controls, and binucleate cells were not detected. However, the cortical NSCs did have a reduced midbody index and wider, disorganized midbodies. Together these data suggested that loss of Kif20b specifically causes defects in abscission, rather than in mitosis or furrowing. Furthermore, NSC apoptosis was elevated in the mutant embryo brains, suggesting that abscission defects cause apoptosis in a fraction of NSCs (26).

Here we address two main questions: whether cytokinesis is differentially regulated in NSCs during earlier and later stages of cortical development, and whether loss of Kif20b disrupts the kinetics of furrowing or abscission. To do this, we developed methods to quantitatively analyze cytokinetic furrow ingression and abscission in NSCs through live imaging in intact cortical explants, and compared different developmental stages when divisions are predominantly proliferative or neurogenic. Our data suggest that abscission in NSCs is usually bilateral, consistent with the idea that the MBR is released at the ventricular surface, and has cell to cell signaling potential. Interestingly, MBRs are observed more often with proliferative divisions than neurogenic divisions. Furthermore, *Kif20b* appears to regulate both furrow ingression speed and abscission timing in cortical NSCs. Together these data add to a growing body of work showing that subtle alterations of abscission timing and disposal of MBRs can influence development.

## Results

### Furrowing and abscission kinetics of embryonic NSCs can be analyzed in intact cortical explants

To better understand cytokinesis events and kinetics of NSCs in their polarized epithelial structure, and to identify specific defects in cytokinesis mutants, we developed a method to live image both cytokinetic furrowing and abscission in intact cortical explants. Both furrowing and abscission occur near the apical membrane. Therefore, instead of cutting cross section slices, we dissect neocortices from membrane-GFP transgenic mice (Figure 1A), and culture them as flat explant “slabs” in glass-bottom dishes to view the apical membrane *en face*. The cultured slabs, which are ∼150 to 200 µm thick, are incubated with SiR-Tubulin (Silicon-Rhodamine-Tubulin), a cell-permeable dye which fluoresces in far red when it binds polymerized microtubules. Thus we can image fields containing many NSC apical endfeet, from the apical membrane up to 15 µm deep. Cells in mitosis are rounded up with a bipolar spindle, making them easily detectable using membrane-GFP or SiR-Tubulin (Figure 1B, C, D, subapical planes). Midbodies form at the apical membrane (Figure 1 B, C, D, apical planes). Live time-lapse imaging of NSC cytokinesis can be done using time points 3 minutes apart for single color imaging and 15 minutes apart for two-color imaging. These time intervals and short exposure times are both essential for maintaining the health of the NSCs. We live-imaged cytokinesis in both control and *Kif20b* mutant cortical explants at two ages.

### Kif20b may facilitate the speed of polarized furrow ingression in cortical NSCs

NSC cleavage furrowing is essential for setting up the plane of division and positioning the midbody at the apical membrane. Here, we set out to determine the kinetics of furrowing and if they change with developmental stage or loss of Kif20b. Mitotic cells are easy to identify near the apical membrane owing to their large, spherical shape. Cytokinesis begins with the cell elongating parallel to the apical membrane before the furrow initiates on the basal side, and ingresses asymmetrically towards the apical membrane. In control cells, furrow ingression proceeds smoothly, at a constant rate (Figure 2Aa). Under these conditions, the rate of furrow ingression for control NSCs is 0.4 µm/min (Figure 2B, E, black lines) and takes ∼15 minutes to complete (Figure 2D, G, control), at both E11.5 (proliferative divisions) and E13.5 (neurogenic divisions). The elongation is also unchanged during development (Figure 2C, F, black lines). Most *Kif20b -/-* NSCs behave like controls, and none show furrow regression (Figure 2Ab). However, a subset furrow abnormally, taking 24 minutes to complete (Figure 2Ac, B, E, red lines). This subset of slow cells causes the median time to furrow completion to be increased by 3 minutes in *Kif20b* mutant brains. These data show that 1) furrowing kinetics do not significantly change between E11.5 and E13.5, and 2) Kif20b is not required for cleavage furrowing, but may help to ensure continuous rapid furrow ingression.

**Figure 2:**
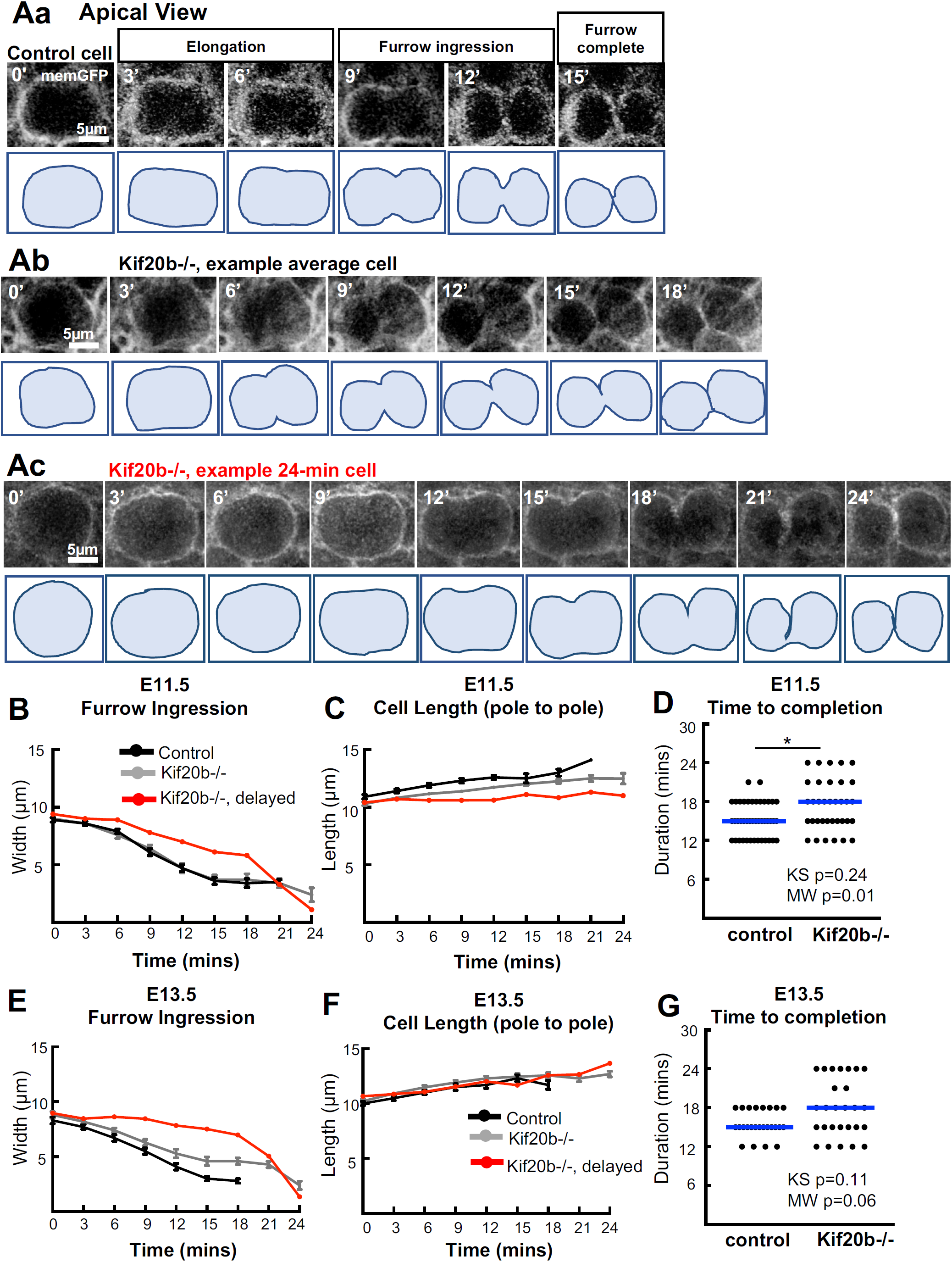
The rate of polarized furrow ingression is regulated by Kif20b. (Aa-Ac) Representative images and traced silhouettes of cytokinetic furrow ingression in one E13.5 control (Aa), one normally furrowing *Kif20b* mutant (-/-) NSCs (Ab, average) and on abnormally furrowing mutant (Ac, 24 min). Furrowing was considered complete when no additional narrowing of furrow width occurred, and the membrane between sister cells appeared continuous. (B-F) Furrow width and pole to pole length were plotted over time in E11.5 (B, C,) and E13.5 (E, F) control and *Kif20b-/-* cortical slab explants. Most *Kif20b-/-* cells furrow at a similar rate to controls, but some furrow more slowly (red line). (D, G) Time to furrow completion was increased in E11.5 and E13.5 *Kif20b* mutant cortices with a subset of NSCs taking 24 minutes to complete (blue lines = medians). E11.5: n=43 control cells from 2 brains (1 +/- and 1 +/+), and 35 *Kif20b-/-* cells from 2 brains. E13.5: n=25 control cells (from 2 +/- and 1 +/+ brains), and 27 *Kif20b-/-* cells from 3 brains. Kolmogorov-Smirnov (KS) and Mann-Whitney (MW) for D, G.

### Abscission is accelerated in Kif20b mutants

It was unknown how long abscission takes following mitosis in cortical NSCs. Therefore, we analyzed the spatial and temporal dynamics of abscission in NSCs in the cortical explant system. Figure 3A shows a representative NSC at an early stage of abscission, with a wide pre-abscission midbody. A rotated image stack (X-Z view) illustrates how the midbody microtubule bundle bridges the two daughter cells across their cell junction. At later stages of abscission (Figure 3B), the midbody is narrower, and sometimes microtubule bundle thinning can be seen on one flank (open arrowhead). By live time-lapse, with two-color imaging of the membrane and microtubules, we were able to follow NSCs through abscission. Abscission involves both the disassembly of microtubules and scission of membrane on the midbody flanks. We cannot detect the midbody membrane, since the membrane-GFP label (myristoylated-GFP) does not enrich in the midbody. Therefore, we defined abscission completion as the time point when microtubules were disassembled on a midbody flank, ascertained when the tubulin signal intensity decreased to background level (Figure 3C). Microtubule thinning prior to complete disassembly was sometimes detected (Figure 3C, 60’).

**Figure 3:**
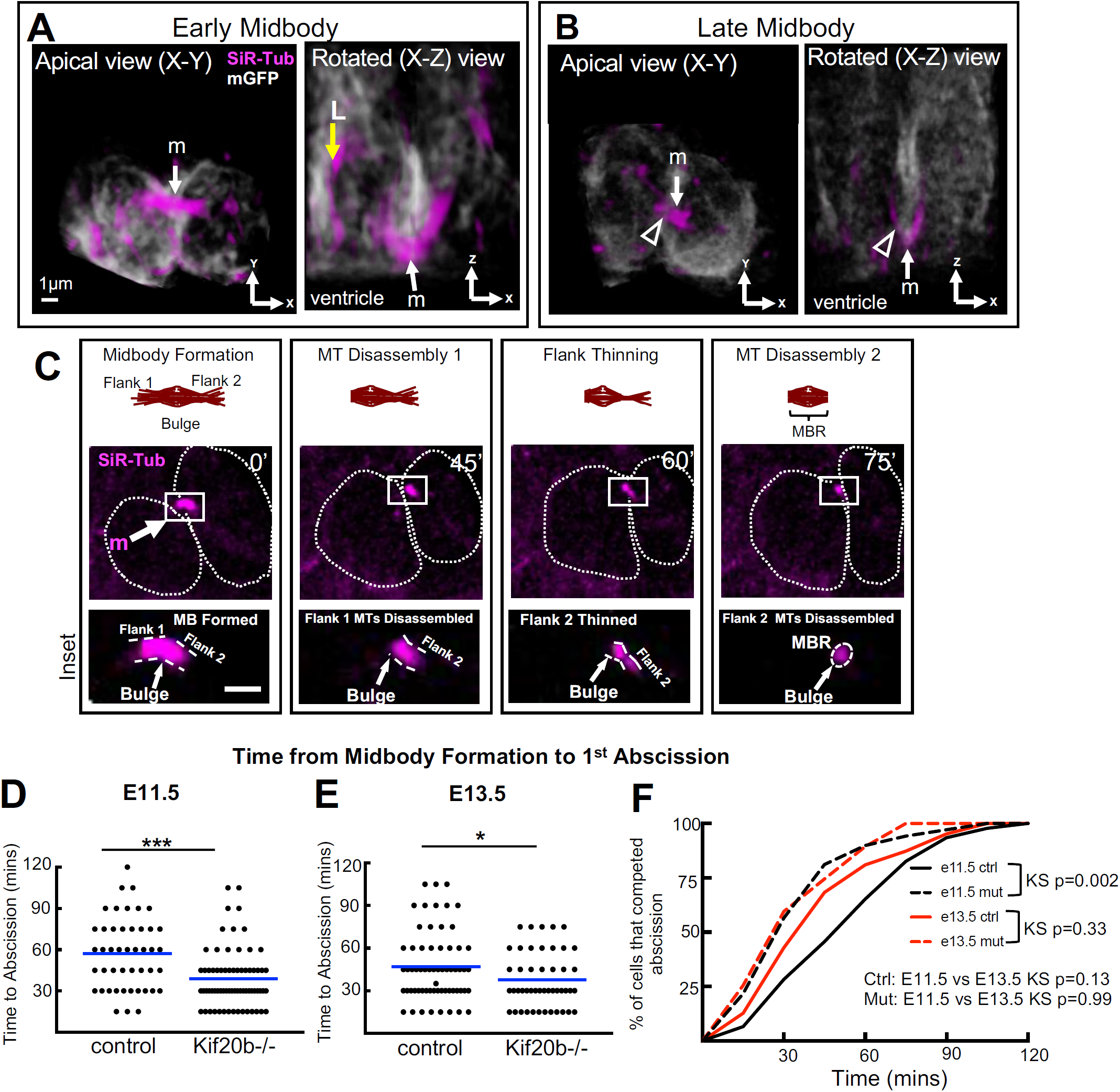
Abscission is accelerated in *Kif20b* mutant cortex. (A, B) 3-D renderings of NSCs at early (A) or late (B) midbody stages labeled by membrane-GFP and SiR-tubulin, viewed from both the apical view (imaging plane) and rotated lateral view. Midbodies (m, arrow) form at the apical membrane (ventricular surface). Late midbody has thinning on one flank (open arrowhead). L, longitudinal microtubule bundles. (C) Schematic and images of time-lapse imaging of an NSC midbody undergoing abscission with bulge and flanks indicated in insets. Distinct steps seen here are: midbody formation (m), microtubule disassembly on flank 1 (1^st^ abscission), flank 2 thinning, and microtubule disassembly on flank 2 (2^nd^ abscission). Dotted circular outlines show shape of sister cell plasma membranes at a subapical plane where cells are the widest. (D, E) Time from midbody formation to 1^st^ abscission is reduced in *Kif20b* -/- NSCs at both ages (E11.5 and E13.5). Blue lines show means. (F) Cumulative frequency plots show *Kif20b -/-* NSCs abscission timing curve is shifted to the left and has an altered shape. For E11.5, n= 46 +/+ cells (from 4 brains); 69 *Kif20b*-/- cells (4 brains). For E13.5, n= 63 +/+ cells (5 brains); 47 -/- cells (3 brains). T-test for D, E. Kolmogorov-Smirnov (KS) and Mann-Whitney (MW) for F.

The timing of abscission has been linked to daughter cell fates (19, 29). To ask if abscission duration changes as brain development proceeds, we measured the time from midbody formation to the first abscission at two different ages. At E11.5, when divisions are predominantly proliferative, we found that abscission duration ranged widely, from 15 to 120 minutes, with a mean of 57 minutes (Figure 3D). At E13.5, when neurogenic divisions have increased, the range of durations was 15 to 105 minutes, and the mean 47 minutes (Figure 3E). A cumulative frequency plot shows a trend to faster abscission at E13.5 (Figure 3F, solid lines). These data suggest that abscission duration may increase as development proceeds. However, we were not able to test this at a later age when divisions are primarily neurogenic, since the thicker cortical slabs from older brains (>400um thick at E15.5) do not remain healthy in these cultures.

Based on our previous findings in fixed brains, that *Kif20b* mutant NSCs have abnormal midbody structure, and increased apoptosis (26, 30), we hypothesized that some *Kif20b* mutant NSCs have delayed or failed abscissions. To our surprise, in live imaging, every mutant NSC observed was able to complete abscission. Moreover, the average time from midbody formation to the first abscission was not delayed, but accelerated compared to controls, by 18 minutes at E11.5, and 9 minutes at E13.5 (Figure 3D, E). Comparing the cumulative frequency plots of *Kif20b* mutant abscissions at the two ages shows they almost overlie (Figure 3F, dashed lines), suggesting a loss of developmental regulation. Thus, Kif20b regulates the duration of abscission, especially in early cortex when symmetric proliferative divisions predominate.

Reports vary on whether abscission occurs on one midbody flank or both flanks, suggesting that it may depend on cell type or context. Therefore, we wanted to determine the frequency of cortical NSCs abscission on one or both midbody flanks, the relative timing, and if this varies with developmental stage or when Kif20b is absent. Interestingly, MBRs have been found in cerebrospinal fluid of early (E10.5) mouse embryos (31), and it was suggested that bilateral abscission is important for symmetric fates in early proliferative NSC divisions. Surprisingly, we detected bilateral abscission just as often in E13.5 divisions as E11.5 divisions, in at least 60% of the cases (Figure 4A, B). The two abscissions usually were captured in sequential images, but occasionally both took place within one time-lapse interval. The median time between the 1^st^ and 2^nd^ event was 30 minutes at E11.5 and 15 minutes at E13.5, with a maximum of 90 minutes (Figure 4C, D). Kif20b loss does not appear to alter the frequency of bilateral abscission (Figure 4 A, B), but it does significantly reduce the time between first and second abscissions in E11.5 NSCs (Figure 4C). In total, the mean time to complete bilateral abscissions was 23 minutes less in the *Kif20b* mutants at E11.5, and 13 minutes less at E13.5 (Figure 4E, F, G). Together these data suggest that bilateral abscission is common in NSCs, and that Kif20b does not regulate its frequency, but regulates the timing of its completion.

**Figure 4:**
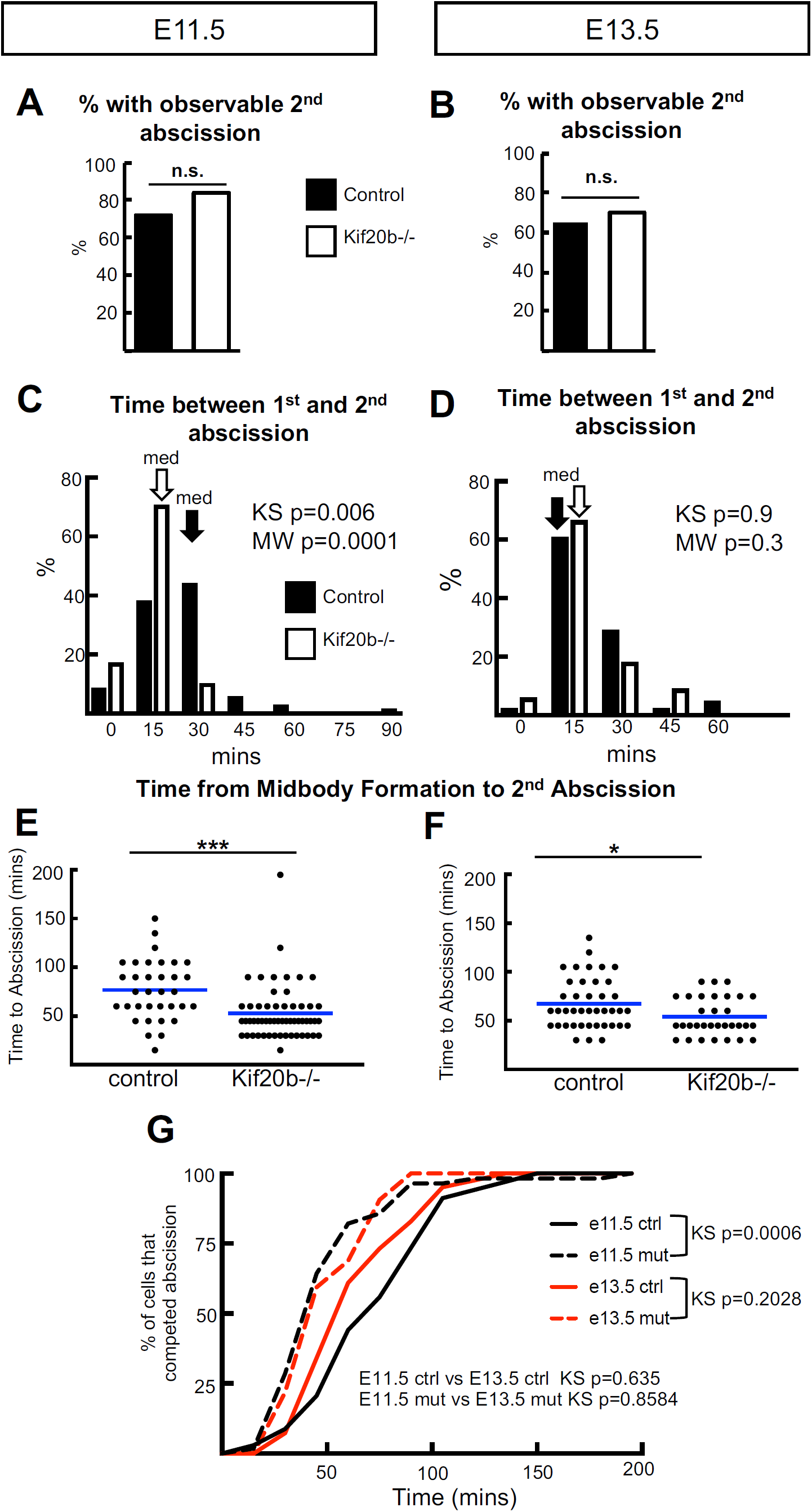
Sequential abscissions on both flanks are observed in most NSC divisions, and occur faster in *Kif20b* -/- brains. (A, B) Using the same methods as in Figure 3, the 2^nd^ abscission on the other flank was observable in more than 60% of cells in E11.5 and E13.5, regardless of genotype. (C, D) The 2^nd^ abscission was usually detected within 30 minutes of the first, but within 15 minutes in E11.5 *Kif20b* -/- cells. (E, F) The total time from midbody formation to completion of bilateral abscissions are reduced in *Kif20b* mutant NSCs at both ages. Blue lines show means. (G) Cumulative frequency curves of 2^nd^ abscissions of E11.5 and E13.5 *Kif20b -/-* NSCs are shifted to the left (dashed lines). For E11.5, n= 34 +/+ (4 brains); 56 *Kif20b-/-* cells (4 brains). For E13.5, n= 41 +/+ (5 brains); 32 *-/-* cells (3 brains). Fisher’s Exact test for A, B. Kolmogorov-Smirnov (KS) and Mann-Whitney (MW) for C, D, G. T-test for E, F.

### Midbody remnants are abundant in early cortex and associated with proliferative NSC divisions

Post-abscission midbody remnants (MBRs) have been shown to adhere to plasma membranes of daughter or neighbor cells, and may influence cell polarity or proliferation (22, 24, 32, 33). We wanted to ask whether MBRs of cortical NSCs remain on the apical membrane after abscission, and whether this changes across development. To do this we first used fixed cortical slabs to assay for MBRs. MBRs can be identified by immunostaining slabs for endogenous Citron kinase (CitK) and Aurora Kinase B (AurKB) (Figure S1, (15)). Both pre-abscission midbodies (brackets) and post-abscission MBRs (arrowheads) are observed at the apical membrane (Figure 5A). Post-abscission MBRs retain the central bulge marked by CitK but lose the AurkB-labeled flanks, and they appear as a disc or donut. We quantified MBRs on the apical surface at two ages, when divisions are primarily proliferative (E11.5) or neurogenic (E15.5). Strikingly, ∼4-fold more MBRs are observed on E11.5 cortices than E15.5 (Figure 5B). Kif20b loss does not alter the frequency of observed MBRs or the developmental difference. These data suggest that E11.5 and E15.5 brains may have different regulation of either MBR release or degradation, but that Kif20b does not regulate these processes.

**Figure 5:**
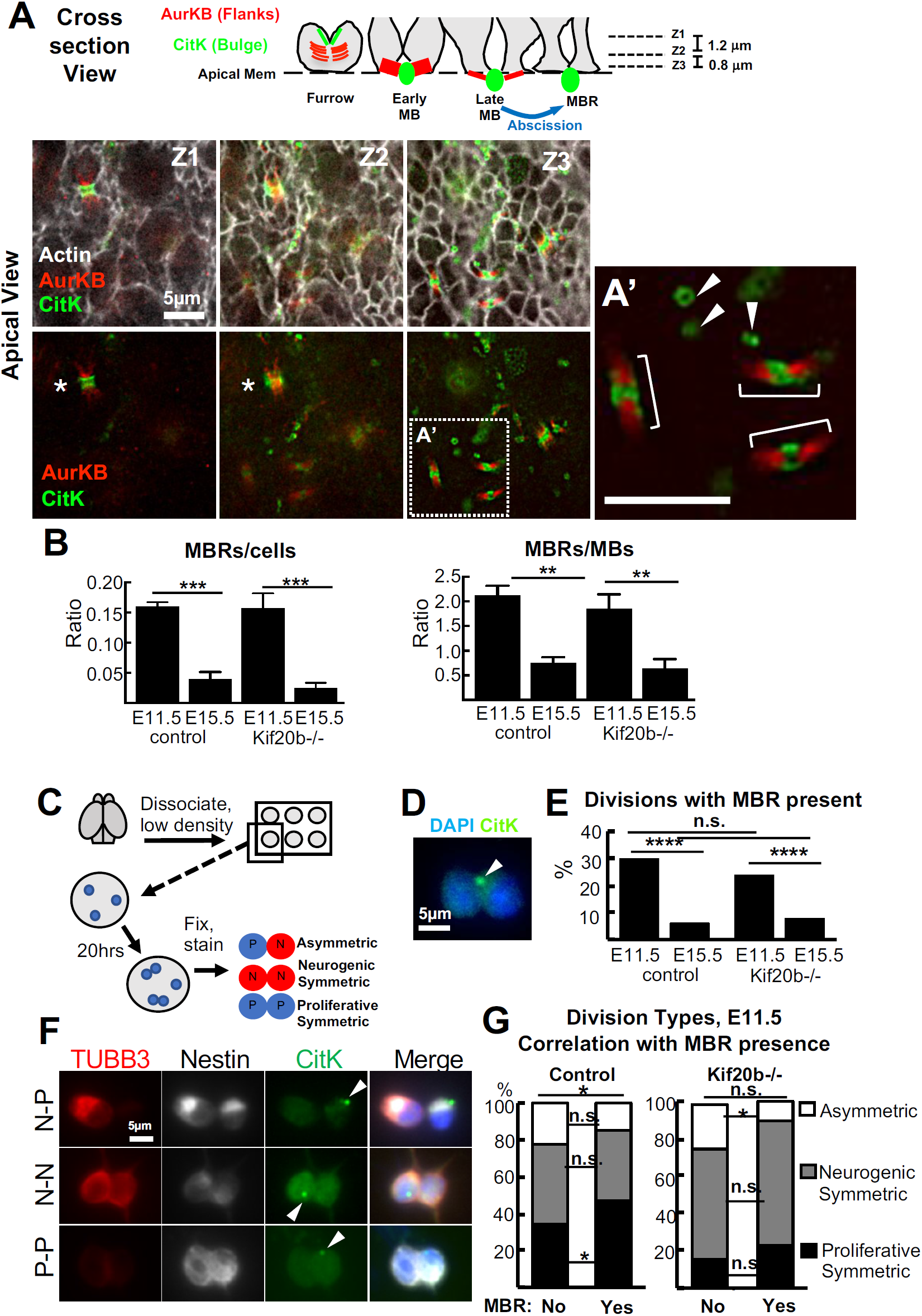
Midbody remnants (MBR) are detected at the apical membrane, and correlated with early proliferative symmetric divisions. (A-A’) Schematic and three image-planes of a field of apical membrane of an E11.5 cortex show many NSC junctions (actin, white), a late furrow in Z1-2 (asterisks), pre-abscission midbodies (brackets, A’), and post-abscission MBRs in Z2-3 (arrowheads, A’). AurkB labels midbody flanks and CitK labels midbody bulge and midbody remnants. (B) Many more MBRs are seen on E11.5 apical membranes than E15.5, normalized to either cell number or midbody number. For E11.5, n= 4 control (3 +/- and 1 +/+); 4 *Kif20b*-/- brains. For E15.5, n= 3 control (1 +/- and 2 +/+); 4 -/- brains. (C) Schematic of pair-cell assay. (D) Pair of daughters of an E11.5 NSC division *in vitro*, with an associated MBR labeled by CitK (arrowhead). (E) E11.5 NSC division pairs are more likely to have a MBR present than E15.5 division pairs. This difference does not depend on Kif20b. (F) Images of three division pairs with a MBR (CitK+) detected. P, progenitor (Nestin+, Tubb3-); N, neuron (Nestin+, Tubb3+). (G) The proportions of division types were compared in pairs with or without a MBR present. Control NSC division pairs with MBRs are more often proliferative symmetric (P-P). *Kif20b* -/- NSCs have reduced proliferative symmetric divisions, with or without a MBR present. For E11.5, n= 394 control (3 +/-, 2 +/+ brains); 272 *Kif20b*-/- divisions (4 brains). For E15.5, n= 241 control (4 +/-, 1 +/+ brains); 280 *Kif20b*-/- divisions (5 brains). T-test in B and the error bars are standard error of the mean. Fisher’s exact test in E, G. Chi-square test in G.

To more precisely quantify MBR association with individual NSC divisions, we modified the established NSC “pair cell” assay (34). NSCs were plated as single cells at clonal density, and fixed the next day to assess whether MBRs are present on newly divided daughter pairs (Figure 5C, D). First, we found that with control NSCs, ∼30% of E11.5 division pairs have a MBR, while only 5% of E15.5 division pairs do, an ∼6-fold difference (Figure 5E*). Kif20b-/-* divisions pairs showed a similar developmental difference. Then, to ask whether MBR were associated with particular daughter fate outcomes, we scored MBR presence in the different types of divisions of E11.5 NSCs. By co-labeling with NSC marker Nestin and neuron marker tubulin-beta-III (Tubb3), the division pairs were classified as proliferative symmetric (2 NSC daughters), neurogenic symmetric (2 neuron daughters), or asymmetric (Figure 5F). Interestingly, E11.5 division pairs that have an associated MBR are more likely to be proliferative symmetric (Figure 5G, control). In the *Kif20b* mutant, this association is no longer statistically significant; however, this may be precluded by a significant reduction in the percentage of divisions that are proliferative symmetric, compared to control NSCs (Figure 5G, *Kif20-/-* bars). The reason for this is unknown, but does not seem to be due to a loss of MBRs. Together with the *in vivo* quantifications of MBRs, these data show that MBRs are more prevalent in the early cortex, and are more associated with early proliferative NSC divisions than later neurogenic divisions. Furthermore, Kif20b loss does not disrupt MBR production or prevalence in the early cortex.

## Discussion

The polarized form of cytokinesis in embryonic cortical NSCs is poorly understood, but may influence the segregation of organelles and apical fate determinants to daughter cells as they make fate choices. Recent findings that mutations in *Kif20b* and other midbody genes cause microcephaly in humans and mice suggest that brain development is especially sensitive to defects in cytokinesis (26, 35-39). To elucidate these issues, we developed methods to quantitatively analyze furrowing and abscission in NSCs of the developing cerebral cortex. Here we addressed 1) whether cytokinesis parameters differ as development proceeds from more proliferative to more neurogenic divisions, and 2) how the loss of kinesin Kif20b affects cytokinesis kinetics in the developing cortex. Since the *Kif20b* mutant is microcephalic, these findings have implications for NSC proliferation. While furrow ingression kinetics did not appear to change with developmental stage, Kif20b loss slowed the rate of polarized furrow ingression in cortical NSCs. Abscission processes in cortical NSCs do appear to be developmentally regulated. Midbody remnants are more prevalent on the apical membranes of early stage cortices, and are more associated with proliferative NSC divisions than neurogenic divisions. Kif20b loss did not disrupt this developmental regulation of MBRs. However, the loss of Kif20b resulted in accelerated abscission. In addition, *Kif20b* mutant NSCs make fewer symmetric proliferative divisions. In the context of emerging data in both the neurogenesis and cytokinesis fields, these data suggest that subtle changes in the kinetics of cytokinesis or in the disposal of midbody remnants could influence NSC daughter fates.

Why would loss of Kif20b from the midbody cause abscission to occur faster on average? The answer is not clear, but prior work suggests two hypotheses. First, previous work by us and others shows that Kif20b can cross-link microtubules *in vitro* (28), and helps organize and stabilize axonal microtubules in neurons (40). Thus when Kif20b is absent, increased instability of midbody microtubules may make it easier for microtubule severing enzymes like spastin to mediate disassembly. Second, we previously identified a role for Kif20b in keeping NSC midbodies aligned with the apical membrane (26, 30). Hypothetically, if Kif20b links midbody microtubules to apical junction proteins, it could mediate tension on the midbody, which has been shown to affect abscission timing in HeLa cells (41). Much more work is needed to understand how Kif20b and other mechanisms regulate abscission timing in developing tissues.

It is possible that altered abscission timing in NSCs could affect daughter fates or proliferation. Our data showed a trend for faster abscission at E13.5 than E11.5. Consistent with this, in an ES cell line, faster abscission was correlated with more differentiation and reduced potency (29). Interestingly, delayed abscission timing in the fly germline is important for maintaining stem cell fate (19). In HeLa cells, delayed abscission causes persistent intercellular bridges, midbody regression/binucleation, or cell death (42), but faster abscission had no morphological or fate consequences (43). In addition, *Kif20b* mutant NSCs have a significant reduction in proliferative symmetric divisions at E11, about half the control percentage. This is accompanied by a concomitant increase in neurogenic divisions. It is tempting to speculate that the faster abscission in the *Kif20b* mutant contributes to the reduction in proliferative symmetric divisions, and thus to a smaller stem cell pool, and resulting microcephaly. However, we cannot rule out some other role of *Kif20b* in promoting symmetric proliferative divisions.

The number of midbody remnants at the apical membrane is much higher in early stage cortices, even when normalized for number of divisions. The mechanism for this developmental difference is not known. But the correlation of MBR presence with proliferative symmetric fates is intriguing, because MBRs have been proposed to have signaling roles and to contain fate determinants (24). An attractive hypothesis is that an early symmetric proliferative division of a cortical NSC would occur through bilateral abscission and release of the MBR extracellularly, whereas a later asymmetric neurogenic division could result from unilateral abscission on one flank and midbody inheritance by the other daughter, promoting asymmetric fates. However, our data argue against this possibility by showing bilateral abscission is observed with similar frequency at two different developmental time points. Another possibility is that the disposal of MBR is regulated differently by E11.5 and E15.5 NSCs. It is not known if NSCs have similar mechanisms for MBR engulfment and degradation as other cell types (21, 22), but differential regulation of these processes in early versus late-stage stem cells could alter the ability of MBRs to influence the polarity or fates of progeny. The composition and signaling capacity of MBRs in stem cells and other dividing cell types require further investigation.

Subtle changes in early brain development can have larger consequences over time. In cortical NSCs, fate-signaling events at the apical membrane happen concurrently with abscission, and could be affected by abscission duration. These include new apical junction building, notch signaling across it, and centrosome docking/ciliogenesis. As these events occur, the MBR could accumulate fate determinants to sequester or transmit. Future studies in multiple stem cell types and developing tissue systems are needed to further elucidate how developmental regulation of cytokinesis contributes to stemness, differentiation, and building tissues and organs.

## Materials and Methods

### Mice

The animal protocol was approved by the UVA IACUC and the colonies maintained in accordance with NIH guidelines. The morning of the vaginal plug was considered E0.5. All embryos were harvested by cesarean section and littermate controls were used when possible. The *Kif20b*^*magoo*^ (*Kif20b-/-)* mutant mouse was generated in an ENU screen (25). It carries a loss-of-function splice site mutation that reduces the protein to undetectable levels (26). The *Kif20b* mutant mouse was crossed with both the mT/mG reporter line (Jax stock #007576) and Sox2-Cre mice (JAX stock #008454) (44, 45) to produce mice that expressed plasma membrane-localized GFP.

### Antibodies and immunofluorescence staining

Following fixation, cortical slabs or NSC cultures were incubated for an hour at room temperature (20 °C) in blocking buffer (0.1% Triton-X, 2-5% Normal Goat Serum in PBS) and then overnight at 4 °C or for three hours at room temperature (20 °C) in the appropriate primary antibody solution (antibody diluted in blocking solution). After primary incubation, coverslips (Fisher Brand Microscope Coverglass 18Cir-1D Cat #: 12-545-84) were rinsed in PBS (3 times every 10 minutes) or slabs were rinsed in PBS-triton once followed by two PBS rinses and then incubated at room temperature (20 °C) with appropriate secondary antibody solution (1:200 dilution) for 30 minutes (cultures) or 45 minutes (slabs) in the dark. Following final washes in PBS, coverslips were mounted onto glass slides with flurogel (Electron Microscopy Sciences 17985-10) or slabs were cover-slipped (22 × 22 #1 cover glass, Curtin Matheson Scientific Inc Cat #: 089-409) with Fluoromount (Vectashield, 4-1000). Antibodies used: chicken anti-Nestin (Aves Lab, NES), rabbit or mouse anti-Tubb3 (TUJ1, Abcam rabbit 52623, Biolegend mouse 801201), mouse anti-Citron kinase (CITK, BD Biosciences 611376), rabbit anti-Aurora B (AurKB, Abcam ab2254), mouse anti-Cep55 (Santa Cruz sc-377018), rabbit anti-Survivin (Cell Signaling Technology 2808). For F-actin visualization we used Phalloidin (ThermoFisher A12380).

### Pair cell assay for NSC divisions

The pair cell assay for dissociated cortical NSC divisions was adapted from (34, 46). E11.5 or E15.5 mouse embryos were dissected from the dam and placed in hibernation medium (Gibco Hibernate-E A12476-01). Cortices were removed and meninges were removed from E15.5 cortices. The cortices were then placed into a 15 mL conical tube for dissociation. Papain from Worthington kit (Worthington Biochemical Corporation, LK003150) was added and cortices were placed on a rotator at 37 °C for 30 minutes (E11.5) or 45 minutes (E15.5). The cortices were then manually dissociated followed by centrifugation at 4 °C at 1300 rpm for 10 minutes. The cell pellet was then washed with DMEM (Invitrogen 11960-051). The wash and centrifugation steps were repeated three times. After the final wash, the cortices were resuspended in final culture medium with a 1 mL pipette followed by a glass Pasteur pipette (resuspension volume varied by age and size of cortices). The cell suspensions were at room temperature (20 °C) for 15 minutes to allow for the clumps to settle to the bottom of the conical tube. Between 1 to 3 µL of cell suspension from the top of the tube were added to each well of the poly-L-lysine coated Terasaki plates (Fisher cat # 07-000-401) to get a very low density. The plates were placed in a humidifying chamber and into a 37 °C incubator with 5% CO_2_ for 20 hours. Final media (made fresh day of culture): 2 mL100X Na-Pyruvate (Invitrogen 11360–070), 2 mL 200 nM L-glutamine (Gibco A2916801), 4 mL 50X B-27 without Vitamin B (Gibco 12587010) per 200 mL of DMEM (Invitrogen 11960-051) filtered through a 0.22 µm cellulose acetate membrane (BD Biosciences 302995). Once filtered 100 µL of 100X N2 (Invitrogen Cat. No. 17502-048), 100 µL 100X N-acetyl-cysteine (Sigma Cat No A-9165), and 20 µL of 10 µg/mL bFGF (Invitrogen Cat. No. 13256-029) were added per 20 mL of media. The cells were fixed with 4% paraformaldehyde (PFA) for 2 minutes followed by 5 minutes cold methanol (100% methanol at −20 °C) in −20 °C freezer. Immunostaining was followed as described above except the plate was kept in a humidified chamber to prevent evaporation and volumes of washes and antibodies were adjusted. Images were acquired using a Zeiss AxioZoom Observer.Z1 with a 40x air objective.

### Cortical slabs for fixed tissue

This method was previously described (47). Briefly, the skulls were opened to reveal the cortices and the head was fixed for 20 minutes in 2% PFA for E11.5 and 4% PFA for E15.5. Cortical hemispheres were pinched off, placed in slide wells (Cat # 70366-12, 70366-13), and trimmed to flat slabs. The total slab thickness varied by age (E11.5: ∼150µm, E15.5: ∼400µm). The slabs were briefly fixed again with PFA before immunostaining. After coverslipping, images were acquired using the 60x objective on the Applied Precision (GE) DeltaVision microscope.

### Cortical slab explant culture for cleavage furrow and abscission live imaging

E11.5 or E13.5 embryos were removed from the uterus and dissected in cold 1x PBS. After decapitation, the skull was removed, and tweezers were used to pinch out the cortices. Each hemisphere was placed into a glass bottom dish (MatTek P35G-1.0-20-C) containing 50 nM silicone-rhodamine tubulin (SirTubulin) (Cytoskeleton CY-SC002) in final culture media (described above). The hemispheres were then trimmed to create flat slabs and flipped, so the apical surface faced the glass. Each dish was placed in a humidifying chamber and into a 37 °C incubator with 5% CO_2_ overnight (Approximately 15 hours). The next day the cortices were removed from the incubator, and the 50nM Sir-Tubulin-containing media was removed. Matrigel (Corning 356237) (1:3 dilution in final culture media) was added on top of the cortical slab. High vacuum grease (Fisher 14-635-5D) was placed on the edge of the glass bottom dish in three spots, and a coverslip (Fisher 12-545-100) was placed over the top of the slab. Gentle pressure was applied to the coverslip at the spots where the vacuum grease was placed. The dish was put back into the incubator, and the Matrigel was allowed to solidify for 5 minutes. Final culture media with 2% Hepes (Gibco 15630080) was added to the dish, and then imaging was performed. An Applied Precision (GE) DeltaVision with Truelight Deconvolution and softWorx suite 5.5 image acquisition software equipped with a heating plate and 40X objective (1.53 N.A.) was used for time-lapse image acquisition. Z-stack images were taken every 15 minutes for abscission and every 3 minutes for cleavage furrowing for up to 6 hours. To minimize phototoxicity, the neutral density filter was set at 32% or lower and the exposure time was kept to a minimum (< 0.1 milliseconds per slice). Z steps were 0.4 µm, and approximately 10 µm deep into the cortical slab was imaged. The total slab thickness varied by age (E11.5: ∼150µm, E13.5: ∼200µm). To confirm 50nM SiR-Tubulin treatment did not cause mitotic arrest, we fixed and stained for phospho-histone 3, a mitotic marker. There was no difference in the mitotic index post SiR-Tubulin treatment. Additionally, in time-lapse movies the percentage of cells that failed to complete furrowing or abscission was similar between genotypes.

### Cleavage furrow ingression and abscission analysis

Deconvolved full-z-stacks and maximum intensity projection images were analyzed using ImageJ. For cleavage furrowing, membrane-GFP cells were identified in mitosis by their characteristic round shape. Time zero was considered the last time point before furrowing begins. The length and width of the cell were measured until furrow completion. Furrowing was considered complete when (1) there was no additional subsequent narrowing of the furrow tips and (2) the membrane between sister cells appeared continuous. For abscission, time zero was midbody formation as ascertained by SiR-Tubulin appearance. Abscission was determined as the time point where there was complete removal of microtubules on one flank ascertained when the SiR-tubulin signal intensity decreased to background level. Midbody membrane scission was shown to be temporally coincident with microtubule disassembly by several previous publications using DIC or phase imaging in cell lines (41, 48, 49), but here we cannot rule out the possibility that the midbody plasma membrane might remain connected for some period of time after the microtubules are gone. Still image stacks for 3-D renderings (Figure 3A, B) were acquired using longer exposure times and finer z-steps on E13.5 cortical slabs incubated in higher concentration of SiR-Tubulin (200 nM). 3-D renderings were created in PerkinElmer Volocity 3D Image Analysis Software (access to Volocity kindly provided by Dr. Barry Hinton).

### Statistical analysis

Statistical analyses were performed with Excel (Microsoft) and GraphPad Prism. All error bars are standard error of the mean (s.e.m.). Data were tested for normality.

## Online supplement

Supplemental Figure S1 shows that citron kinase localizes to the midbody bulge and remnants along with Cep55.

## Acknowledgments

This work was supported by NIH (R01 NS076640 and R21 NS106162 to NDD) and the Robert R. Wagner Fellowship to KCM. We thank Jessica Neville Little, Xiaowei Lu, Bettina Winckler, Ann Sutherland, Sarah Siegrist, Jung-Bum Shin, Maria Lehtinen, Anthony Lamantia and their labs for advice and discussion. We thank Michael Fleming for Figure 1G images.

## Author Contributions

KCM conceptualized and performed experiments, curated data for Figure 1-5 and Figure S1, wrote the first draft, and edited the manuscript. NDD conceptualized and supervised experiments, and edited the manuscript.

The authors declare no completing interests.

---

**Supplemental Figure 1:**
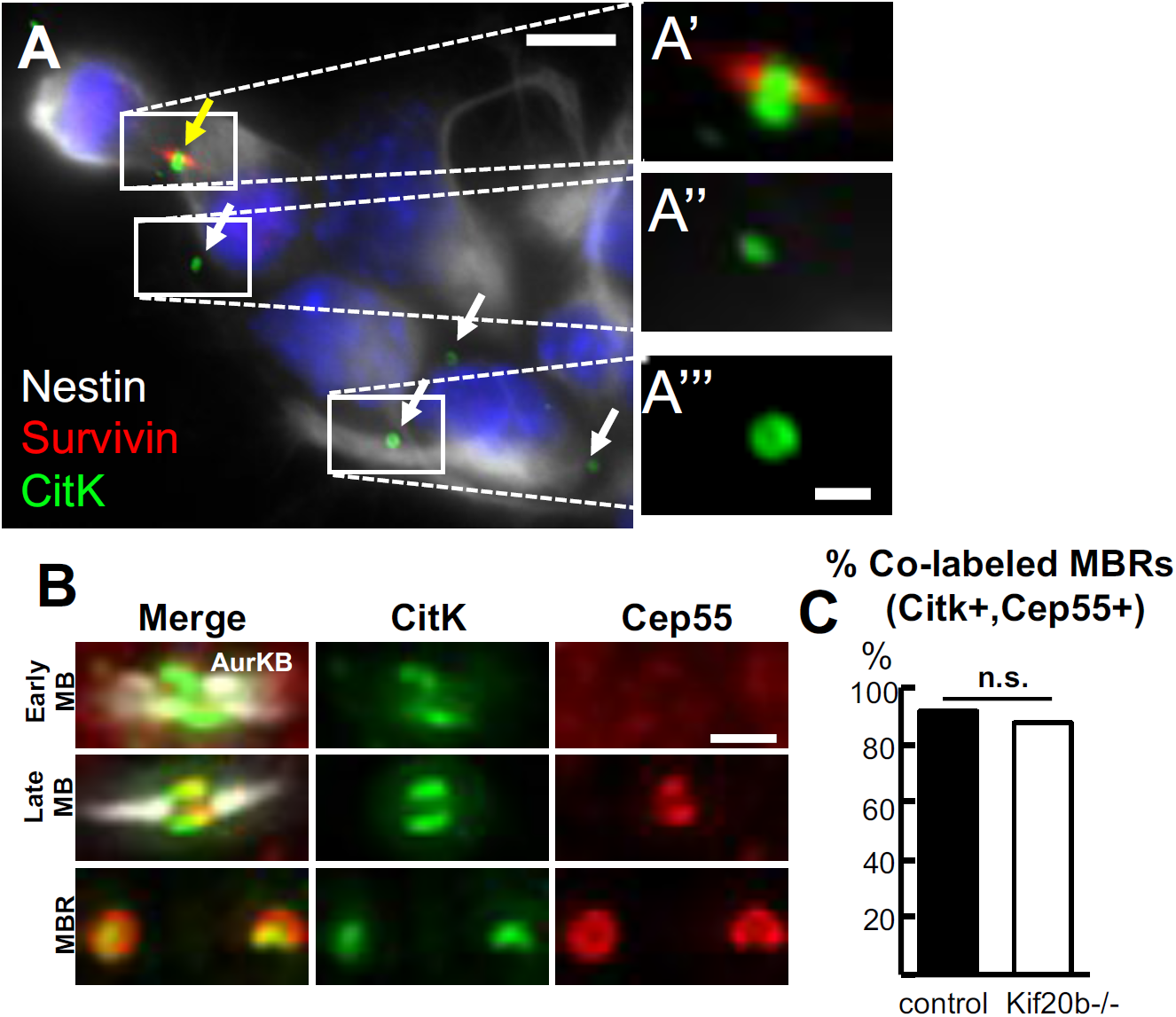
NSC midbody remnants (MBRs) are marked by Citron Kinase (CitK) and Cep55. (A) Dissociated E12.5 NSCs identified by endogenous staining for Nestin. Survivin labeling of midbody flanks shows two daughter cells still connected by a pre-abscission midbody (yellow arrow). CitK appears as a ring around the bulge of midbodies(A’) and MBRs (white arrows, A’’, A’’’(15, 50)). (B) Images of dissociated E11.5 NSCs immunostained for endogenous CitK, Cep55, and AurKB show their localizations at different midbody maturation stages. AurKB localizes to the flanks of early and late midbodies, but is mostly absent in post-abscission MBRs. CitK localizes as a ring around the bulge of early and late midbodies as well as MBRs. Cep55 is absent in early midbodies, but accumulates in a ring around the midbody bulge, but only at late stages, and remains in MBRs (20, 22, 51). (C) Most MBRs are co-labeled with both Cep55 and Citk. Scalebars: 5 µm in A; 1 µm in A’, A””, and B. n= 61 control MBR (1 brain), 49 *Kif20b-/-*MBR (1 brain) n.s: Fisher’s exact test

